# Long-term effects of the early life environment on alternative splicing

**DOI:** 10.64898/2025.11.29.690908

**Authors:** Rose MH Driscoll, Jennifer A Brisson

**Affiliations:** Department of Biology, University of Rochester, Rochester, NY, United States of America

**Keywords:** phenotypic plasticity, polyphenism, early life stress, alternative splicing

## Abstract

All organisms respond to their environments, and early life environments may be especially impactful. Alternative splicing has been implicated as one mechanism that may underlie the response of organisms to their environments. However, most studies of this type examine immediate effects of an environmental perturbation on patterns of alternative splicing; little is known about the long-term consequences of early life environments. Here, we show that the early life environment exerts a lasting influence on patterns of alternative splicing in the pea aphid (*Acyrthosiphon pisum*). This species is sensitive to conspecific density (crowding), with a transgenerational (mother-to-daughter) wing polyphenism cued by this environmental signal. We demonstrate that the maternal environment during embryonic development alters daughters’ alternative splicing profiles independently of morph identity, and that these effects persist into adulthood. By comparing the magnitude of maternal-environment effects to morph-specific differences, we reveal that the influence of early environmental conditions on splicing is unexpectedly large. These findings suggest that environmental history can leave a persistent molecular imprint beyond its role in morph determination. More broadly, this work highlights the need to disentangle environmental and morph-specific transcriptomic variation in studies of developmental polyphenism and connects to broader efforts, across biology and medicine, to understand how early life experiences shape long-term molecular and physiological outcomes.

## 1. Introduction

Phenotypic plasticity encompasses the diverse ways individual organisms respond to their environments (Scheiner, 1993; West-Eberhard, 2003). These responses range from short-term, reversible changes (e.g., Gabriel, Luttbeg, Sih, & Tollrian, 2005) to polyphenism, a distinct form of plasticity in which two or more discrete phenotypes are produced (e.g., Simpson, Sword, & Lo, 2011; Yang & Pospisilik, 2019). Although phenotypic plasticity as a phenomenon has long been recognized, the rise of modern molecular biology has focused attention on the molecular mechanisms underlying these responses (e.g., Aubin-Horth & Renn, 2009; Cossins, Fraser, Hughes, & Gracey, 2006; Sommer, 2020) and, in particular, the ways transcriptomes respond to environmental perturbations. Notably, the transcriptome itself is both a first-order phenotype, and a mechanism that drives higher-order phenotypic changes (e.g., Leung, Grulois, & Chevin, 2022; Renn & Schumer, 2013; Sun et al., 2022).

Transcriptomes can respond plastically to environmental conditions not only through changes in gene expression levels (e.g., Dal Santo et al., 2013; Dang et al., 2021; Vellichirammal, Madayiputhiya, & Brisson, 2016; to name just a few representative studies), but also in terms of the mature mRNAs and resulting proteins produced via alternative splicing (Marden, 2008; Singh & Ahi, 2022). Existing work on this topic has generally focused on two temporal scales of perturbation. The first examines the immediate effect of acute environmental stress, in which individuals are briefly exposed to extreme environmental cues and then changes in splicing are immediately measured (e.g., Healy & Schulte, 2019; Long et al., 2013; Suresh, Crease, Cristescu, & Chain, 2020; Tan et al., 2019; Vitulo et al., 2014; Zhang, Zhang, Yuan, & Li, 2023). The second considers chronic, long-term exposure, in which individuals are reared from early life to adulthood in set environmental conditions for one or more generations (e.g., Choi, Malik, Han, & Kim, 2024; Jaksic & Schlötterer, 2016; James et al., 2012 assesses both timelines). Notably, neither approach isolates the specific influence of early life environments, despite their potential to produce persistent phenotypic effects.

One exception to this general pattern is the study of alternative splicing in environmentally-determined, polyphenic morphs. Such differences have primarily been examined in insects, including alternate wing and reproductive morphs of aphids (Grantham & Brisson, 2018), seasonal morphs of butterflies (Steward, de Jong, Oostra, & Wheat, 2022; Steward, Pruisscher, Roberts, & Wheat, 2024; Tian & Monteiro, 2022), and castes in bees (Aamodt, 2008; Foret et al., 2012; He et al., 2022; Price et al., 2018). Almost by definition, however, polyphenic systems involve coordinated morph development pathways in which external cues toggle a “switch” mechanism that directs individuals down distinct developmental trajectories. Consequently, adult differences (including in splicing) are generally interpreted as outcomes of these coordinated developmental programs, rather than as a direct response to the environment. This distinction is further reinforced by the fact that most studies examine only a single morph per environment, whether by design or convention. As a result, the key question of whether early life environments can exert universal effects (*i.e*., regardless of morph) on patterns of alternative splicing later in life remains untested by these studies.

The wing polyphenism of the pea aphid, *Acyrthosiphon pisum* (Harris 1776), presents a powerful system for disentangling the effects of early environmental conditions from those of programmed morph development on alternative splicing. This polyphenism is cued transgenerationally by maternal crowding: mothers experiencing high conspecific density produce more winged offspring, while mothers in uncrowded environments produce more wingless offspring. However, there is not a one-to-one correspondence between morph and environment in this system; a modest number of the non-favored morph are produced in each environment, likely as a bet-hedging strategy (Grantham, Antonio, O’Neil, Zhan, & Brisson, 2016). Moreover, because daughter embryos develop within their mother’s ovaries, environmental perturbations unrelated to morph determination may be transmitted along with the morph-determining cue. Consequently, by comparing individuals across morph and maternal-environment categories, it becomes possible to distinguish the programmed effects of morph identity from the direct impacts of early life (maternal-embryonic) environments on alternative splicing.

In this study, we demonstrate that the maternal environment exerts a long-lasting effect on daughters’ patterns of alternative splicing. We examine adult pea aphids of both winged and wingless morphs that developed as embryos inside mothers experiencing crowded or uncrowded conditions, but were reared under common garden conditions after birth. Using a two-by-two factorial design, we disentangle the programmed effects of morph identity from the direct impacts of the maternal environment. We compare the relative magnitude and properties of these two classes of effects to reveal the significant contribution of early life environmental perturbations.

## 2. Methods

### 2.1. Aphid populations and maternal treatments

Pea aphid populations and RNA samples were generated as reported in Gregory, McDonough, Frankino, and Brisson (2026). Briefly, a population of pea aphids was started from a single asexual female from line 584. After population expansion, aphids were reared at low density (3-4 aphids per plant on *Vicia faba* seedlings) for at least three generations. Adults were assigned at reproductive maturity (when they began to produce offspring) to either a low density (uncrowded; remaining on low-density plants) or high density (crowded; 12 aphids in a 30 mm Petri dish for 24 hours) maternal environment treatment. The crowding dishes were set up in triplicate (three dishes of 12 aphid mothers) on the same day and the same number of aphid mothers (36) were used for the low density treatment a few weeks later. After treatment, all adults were placed on leaf plates (one aphid in a 100 mm Petri dish containing a *Vicia faba* leaf embedded in agar) for 24 hours for offspring production.

Offspring (the focal individuals of our study) were tracked until adulthood, when they could be visually distinguished as W or WL. On the third day of reproductive maturity, ovaries were dissected out (to remove transgenerational signal) and each individual carcass of a W or WL daughter of a crowded or uncrowded mother was flash frozen with liquid nitrogen, pulverized, and stored at −80 °C in TRIzol (Invitrogen). RNA was extracted with a TRIzol-chloroform method with DNase treatment and RNA Clean & Concentrator-5 kit (Zymo) according to manufacturer’s protocol.

### 2.2. RNA sequencing and read handling

Library preparation and sequencing was conducted by Genewiz (South Plainfield, NJ, USA) to generate Illumina HiSeq 150 bp paired-end reads. Read trimming was performed with Trimmomatic (v0.39) (Bolger, Lohse, & Usadel, 2014) and quality check with MultiQC (v1.14) (Ewels, Magnusson, Lundin, & Käller, 2016). Reads were mapped to the v3.0 pea aphid genome with HISAT2 (v2.2.1) (Kim, Paggi, Park, Bennett, & Salzberg, 2019) and sorted and indexed with samtools (v1.17) (Danecek et al., 2021) and bowtie2 (v2.5.1) (Langmead & Salzberg, 2012). The same bamfiles of mapped reads were subsequently used in both the alternative splicing analysis and differential expression analysis.

### 2.3. Alternative splicing analysis

We examined alternative splicing using rMATS-turbo (v4.1.1) (Y. Y. Wang et al., 2024). Candidate alternative splicing events were inferred from annotated transcripts and exons in the genome annotation file (GTF), including (1) known splice junctions found in transcripts reported in the GTF and (2) novel splice junctions involving exons reported in the GTF in a combination not represented by any transcript in the GTF. Candidate alternative splicing events included 13,020 skipped exon (SE) events, 1,767 mutually exclusive exons (MXE) events, 673 retained intron (RI) events, 1,373 alternative 3’ start site (A3SS) events, and 1,406 alternative 5’ start site (A5SS) events.

For each event, rMATS detects reads supporting the presence of each of two splice forms - an “inclusion form” and a “skipping form” (Figure S1). An “inclusion level” statistic quantifies the relative abundance of the inclusion and skipping forms and represents the rate of detection of the inclusion form as a proportion between 0 and 1. In our dataset, the magnitude of splicing differences between paired groups tended to be small to moderate for any given event, with differences in inclusion levels between groups as high as 0.667, but never as large as 1, which would indicate that one splice form was exclusively used by one group and the other exclusively used by the other.

We filtered candidate splicing events for a minimum data depth to increase confidence in our calculated inclusion level values and subsequent statistical analyses. We followed Grantham and Brisson (2018) and Rogers, Palmer, and Wright (2021) in filtering for at least 20 reads in at least 50% of the samples (regardless of group) included in a contrast. Since downstream analyses compare across contrasts, we further reduced the events examined to only those that met the read depth requirement in all five of the two-way contrasts. This left 4,713 SE events, 1,427 MXE events, 355 RI events, 794 A3SS events, and 727 A5SS events (Table 1). This filtering method excludes genes that are not expressed or only lowly expressed in one or more sample types (for example, genes with winged-specific expression) even though these genes could exhibit undetected splicing differences in other sample types. Thus, true splicing differences resulting from environment and morph may be even more extensive than what is demonstrated here.

**Table 1.**
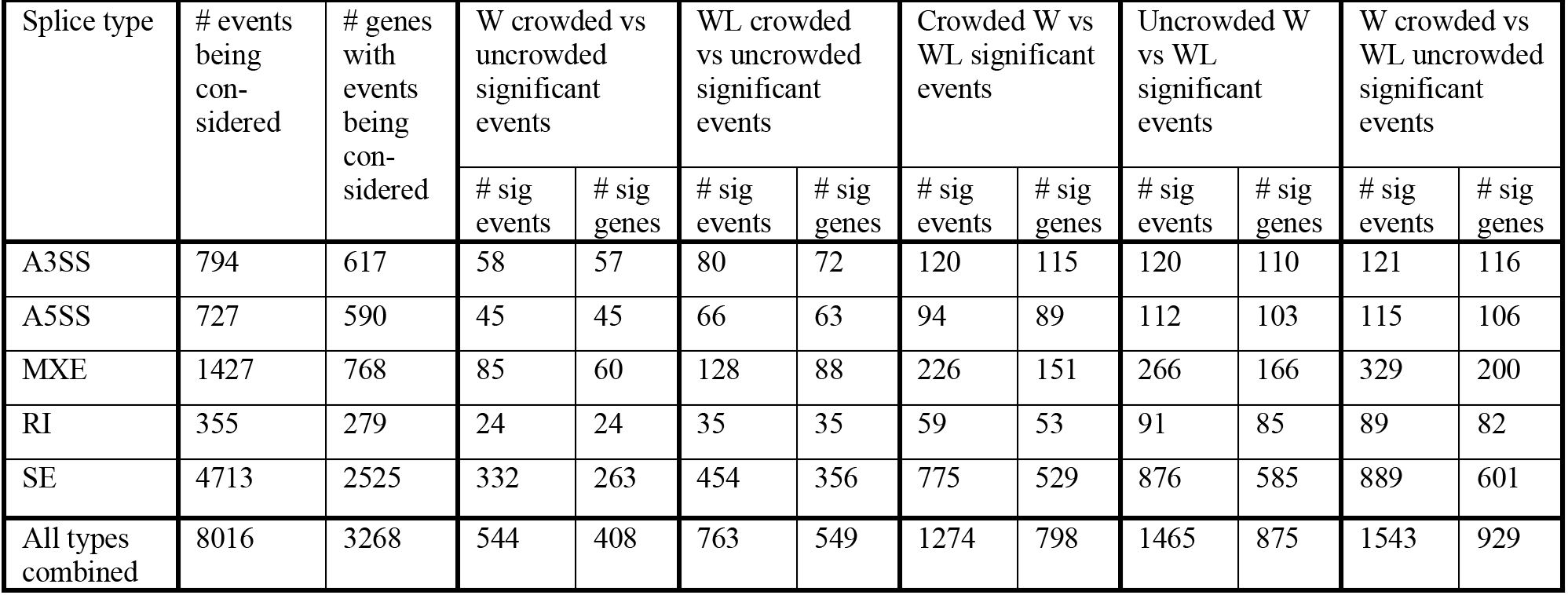
The number of alternative splicing events of each type, and the number of distinct genes those events are found in. W, winged; WL, wingless; crowded, crowded maternal environment; uncrowded, uncrowded maternal environment. A3SS, alternative 3’ start site; A5SS, alternative 5’ start site; MXE, mutually exclusive exons; RI, retained intron; SE, skipped exon.

We investigated dynamics of alternative splicing among these 8,016 events, with the understanding that this does not entirely capture the full transcriptome. To test for significant differences in alternative splicing between two groups, rMATS employs a likelihood-ratio test for differences in inclusion level values. We tested for any difference > 0 by setting rMATS’s -- cstat parameter to 0. In downstream analyses, we did not set any cutoff on the magnitude of inclusion level differences, instead considering all magnitudes of differences as potentially biologically meaningful. As our samples are whole carcasses (minus ovaries) rather than smaller body regions or specific tissue types, a moderate or even small whole-body difference in splice form usage could still correspond to a striking difference in a specific tissue. rMATS is the gold standard program for alternative splicing analysis but, crucially, only supports statistical comparisons between two sample groups. Therefore, we performed our alternative splicing analysis as a series of five paired contrasts between sample groups, followed by an analysis of similarities and differences between paired contrasts sharing particular qualities (e.g., contrasts between different maternal environments, contrasts between different morphs).

This method produces a good understanding of the global patterns of alternative splicing in these samples. To account for the presence of multiple paired contrasts, we discarded the FDR (multiple comparisons correction) values generated by rMATS and manually implemented our own FDR-based multiple comparisons correction to the p-values from all of the tests in the five different paired contrasts all together. Our correction strategy also included all splice types together, although rMATS by default performs FDR correction on each splice type separately.

We conducted the multiple comparisons correction on the collection of all of these p-values in R, using the p.adjust function with method = “fdr” (specifying the Benjamini & Hochberg false discovery rate correction method). We detected significant differences in alternative splicing as FDR < 0.05.

The majority of our analyses involve comparisons of paired sample groups that share one factor (for example, same morph) and differ in the other factor (for example, different maternal environments). As a supplementary analysis, we first combined sample groups with the same maternal environment together in order to compare all crowded versus all uncrowded samples (regardless of their morphs), then conversely combined sample groups with the same morph identity together in order to compare all W versus all WL samples (regardless of their maternal environment). This set of two contrasts was also corrected for multiple comparisons as described above, but this was performed separately from the set of five main contrasts.

### 2.4. Differential expression analysis

Following the example of Rogers et al. (2021), we excluded differentially alternatively spliced exons from the expression quantification, as this can bias estimates. We conservatively removed the exon(s) pertaining to any alternative splicing event that exhibited a significant difference (FDR<0.05) in any paired contrast, without considering data depth (i.e., events which did not meet the required data depth to be included in the alternative splicing analysis could still have their exon(s) excluded from the expression analysis if they had FDR < 0.05). This resulted in the removal of exons involved in 1755 SE events, 538 MXE events, 186 RI events, 346 A3SS events, and 336 A5SS events. For SE events, the single focal exon was removed; for MXE events, the two focal exons were removed; for RI events, the “long” focal exon spanning the two flanking exons and the retained intron was removed; and for A3SS and A5SS events, the longer versions of the focal exons were removed, as the short version represents the constitutively included portion. While this removal of alternatively spliced exons is best practice, a test analysis with these exons included gave broadly similar results, suggesting that alternatively spliced exons do not have a large impact in this dataset.

We obtained read counts for constitutively included exons of each gene with htseq-count (HTSeq v0.9.1) (Putri, Anders, Pyl, Pimanda, & Zanini, 2022). We conducted differential expression analysis with DESeq2 (v1.38.3) (Love, Huber, & Anders, 2014) using a method designed to parallel our alternative splicing analysis workflow as much as possible. Therefore, although DESeq2 has the capability to consider many sample groups at once and test for effects of multiple factors, we instead performed a series of five paired contrasts just as we did for the alternative splicing analysis above, and then proceeded with a comparison of the results of these paired contrasts. We did not apply a log_2_ fold change cutoff, again considering differences of all magnitudes to cast a wide net.

DESeq2 does not report fold change or p-value for genes that had 0 expression in all samples in a contrast. Therefore, in order to compare across contrasts, we removed genes that had 0 expression in all samples in at least one contrast from consideration across all contrasts. We also removed genes that reported count outliers in at least one contrast (according to a Cook’s distance cutoff; DESeq2’s default behavior). This left us with 14534 genes (Table S1). We did not employ DESeq2’s default “independent filtering” behavior (setting independentFiltering to FALSE).

As in the alternative splicing analysis, we discarded the FDR values generated by DESeq2 and manually implemented our own FDR-based multiple comparisons correction to the p-values from all of the tests in the five different paired contrasts all together. We assessed significance at FDR < 0.05.

## 3. Results

### 3.1. The alternative splicing landscape in pea aphids

We quantified alternative splicing in winged (W) and wingless (WL) adult female pea aphids whose mothers were exposed to crowded (high conspecific density) or uncrowded (low conspecific density) environments. We detected 8,016 alternative splicing events which had sufficient data depth for analysis (see Methods). Skipped exons (Figure S1) were the most frequent, with 4,713 events (Table 1; Figure 1B-E). Some genes contained multiple splicing events (Figure S2), such that the 8,016 alternative splicing events are distributed across 3,268 genes (Table 1).

**Figure 1.**
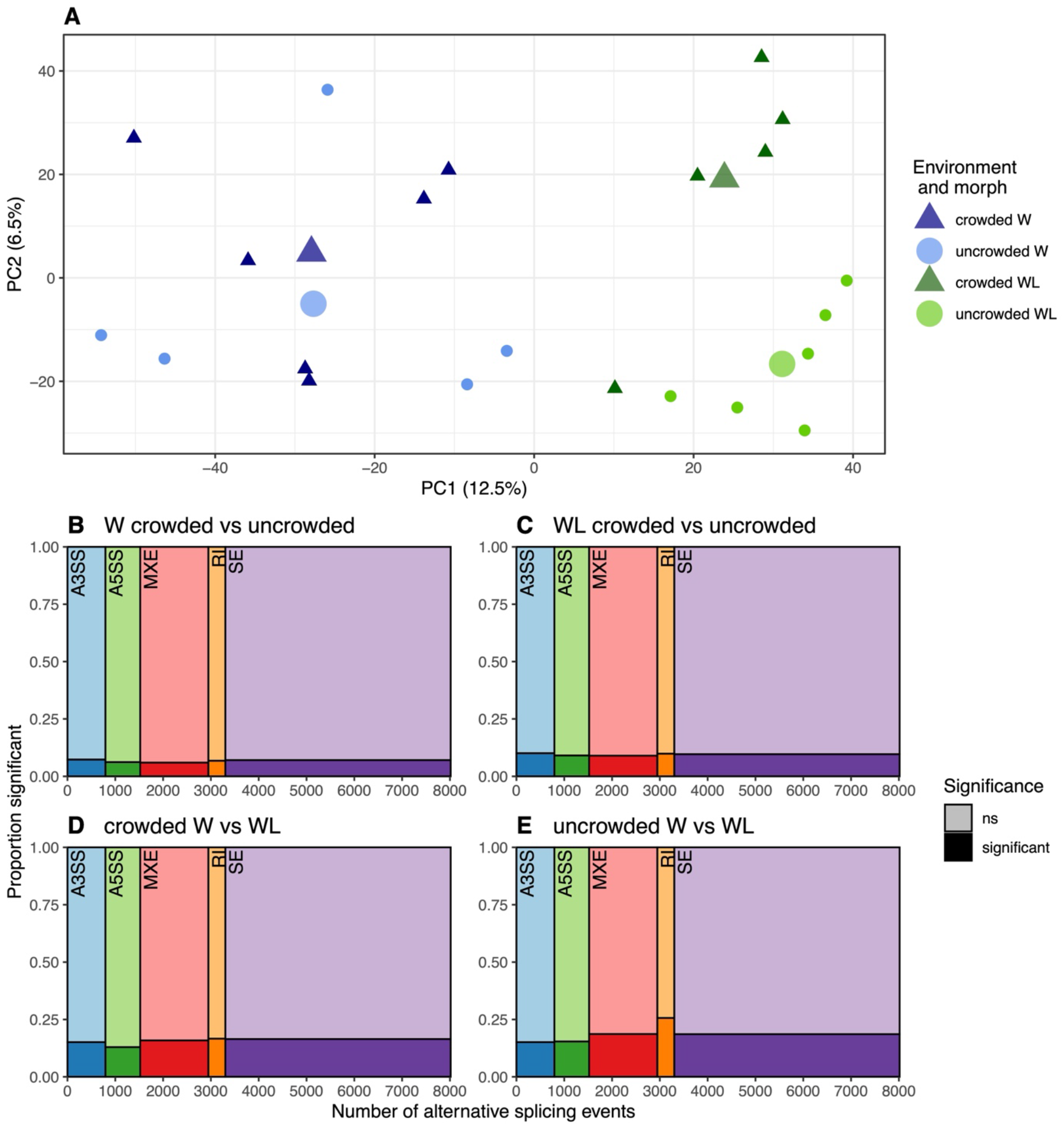
Maternal environment and morph shape the alternative splicing landscape. (A) PCA of per-sample inclusion level values for 8,016 alternative splicing events. Individual samples are represented by small points (winged [W]: blue; wingless [WL]: green; crowded: triangle, dark shade; uncrowded: circle, light shade), while large points represent group means. (B-E) Mosaic plots show the number of significant alternative splicing events of each type in each paired contrast. The x-axis shows the number of total events examined for each splicing type (each represented with a different color), while the y-axis shows the proportion of significant events (FDR < 0.05) in the given contrast (significant: dark shade; nonsignificant: light shade).

### 3.2. Alternative splicing is impacted by both morph and maternal environment

We examined how the maternal environment and morph identity affected alternative splicing. We calculated the relative usage of the two splice forms at a given site in a given sample as a proportion (“inclusion level”) and then used principal component analysis (PCA) to examine patterns across samples (Figure 1A). Morph is the most notable factor in this dataset, with the first principal component (PC1; 12.3% of the variation) separating W and WL samples.

However, effects of maternal environment are also evident, with the second principal component (PC2; 6.6% of variation) separating WL individuals based on maternal environment. We examined each morph separately, and found that with WL samples, the maternal environment effect is fairly coordinated, appearing on PC1 and PC2 ahead of components that represent noise (Figure S3C versus Figure S3D). In W samples, the maternal environment effect is also present but is less coordinated, appearing on PC3 and PC5 while PC1 and PC2 represent noise (Figure S3B versus Figure S3A).

We then tested individual alternative splicing events for significant differences in splicing. A simple comparison between all crowded and all uncrowded individuals (regardless of morph) found 700 splicing events exhibiting significant differential alternative splicing, while a simple comparison between all W and all WL individuals (regardless of environment) found more than twice as many – 1737 significant events. We used a more detailed series of analyses to better understand the specific effects and interactions of these two factors. We made two “environment”-type contrasts, comparing individuals of a single morph from crowded and uncrowded maternal environments, and two “morph”-type contrasts, comparing W and WL individuals from a single maternal environment. We detected 763 significant events (in 549 genes) in the environment-type contrast with WL individuals (Figure 1C; Table 1), and 544 significant events (in 408 genes) with W individuals (Figure 1B; Table 1). Meanwhile, in each of the two morph contrasts, we detected ∼1,400 events (in ∼800 genes) with significant differential alternative splicing (Figure 1D-E; Table 1).

We examined patterns within sets of genes with significantly different splicing events using gene ontology enrichment analysis. Environment-type contrasts generally yielded more enriched gene ontology terms than morph-type contrasts (Table S2), potentially reflecting more functionally cohesive sets of differentially spliced genes in contrasts belonging to the former group, despite their smaller size. However, these results should be interpreted cautiously, as most approach only marginal significance even with p-values not corrected for multiple comparisons. Nevertheless, examining the gene lists (see supplemental files) for each contrast revealed interesting candidates for future study: in the environment-type contrast, these included nutrition-related genes such as insulin receptor substrate (LOC100569479) and putative leptin receptor related gene (LOC100167873); in the morph-type contrast, notable candidates included muscle-related genes [*e.g*., myosin heavy chain (LOC100167379) and troponin (LOC100163274)] and hormone-related genes [*e.g*., ecdysone 20-monooxygenase (LOC100165833) and farnesol dehydrogenase (LOC100160714)].

We conclude that both the early life environment and morph impact alternative splicing. Notably, the number of significant events in the environment-type contrasts is about one-quarter to one-half of the number of significant events in either of the two morph-type contrasts; *i.e.,* the silent reshaping of the alternative splicing transcriptome by the early life environment rivals that of morph differences, with their dramatically different morphologies.

### 3.3. Inconsistent effects of maternal environment and consistent effects of morph

We next assessed the consistency of the alternative splicing responses, reasoning that early life environment treatments may bring about non-directed changes, while morph may involve more concerted changes. We first examined the two maternal-environment-based contrasts (individuals from crowded versus uncrowded maternal environments for each of the two morphs; Figure 2A-D). We found that a relatively modest fraction (W: 20.8%; WL: 14.8%) of the events significant in one environment contrast were also significant in the other environment contrast (Figure 2D), with the majority of significant events being exclusive to one contrast or the other (Figure 2B-C). However, the rate of shared significant events (Figure 2D) among all events was greater than that expected due to chance (*X*^2^ test of independence on a 2×2 contingency table of significance in the W crowded vs W uncrowded contrast versus significance in the WL crowded vs WL uncrowded contrast; p < 2.2×10^-16^). Events that were significant in both maternal environment contrasts had the same direction of effect (positive or negative; Figure 2D, upper right and lower left quadrants) in both contrasts 69.0% of the time, more often than would be expected due to chance (*X*^2^ test of independence on a 2×2 contingency table of direction of effect in the W crowded vs W uncrowded contrast versus direction of effect in the WL crowded vs WL uncrowded contrast; p = 2.00×10^-4^).

**Figure 2.**
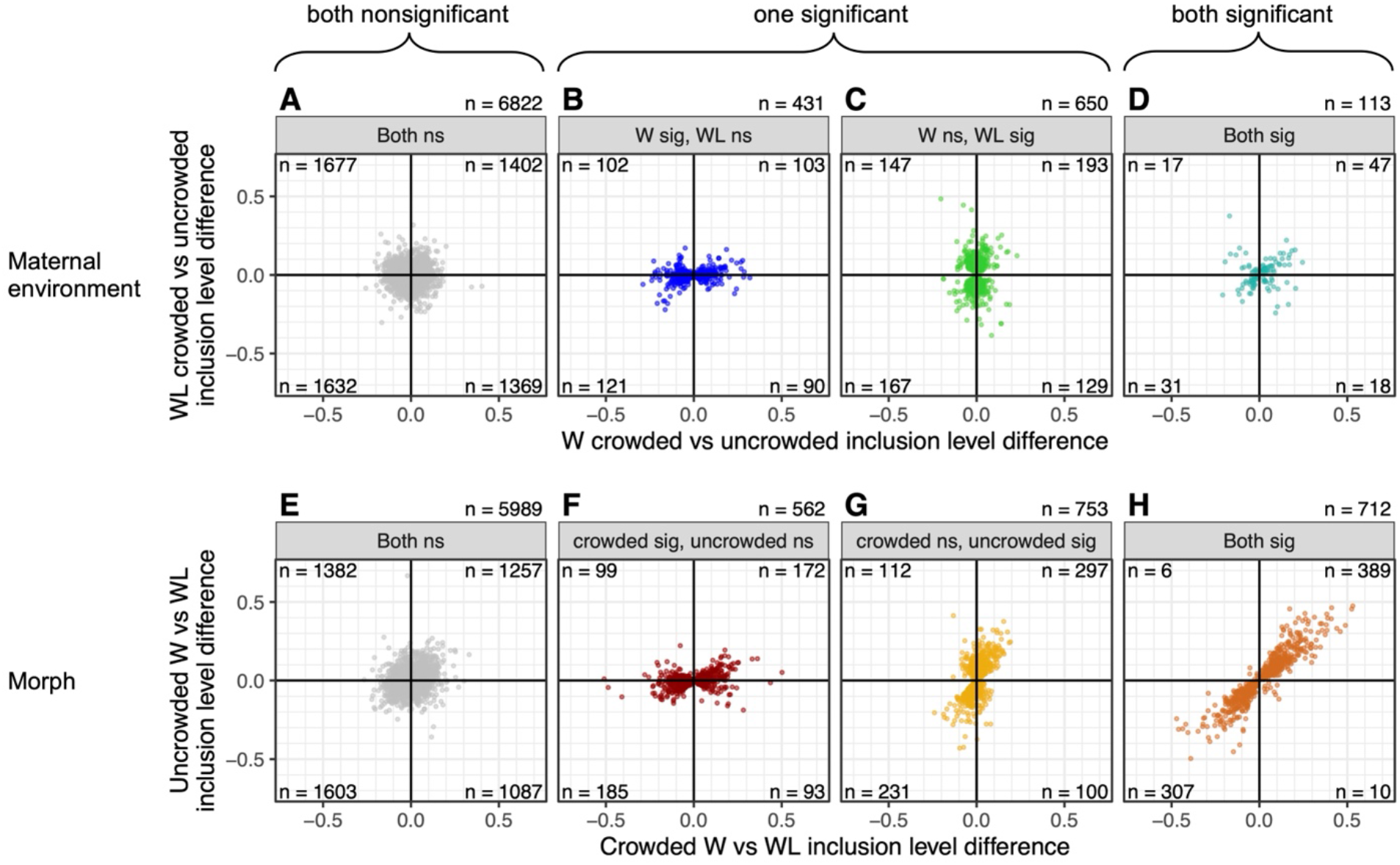
Inconsistent effects of maternal environment and consistent effects of morph. Scatterplots in panels A-D show the difference in inclusion level for each alternative splicing event between individuals from a crowded versus an uncrowded maternal environment for W (on the x-axis) versus WL (on the y-axis) for four significance categories: (A) events that were nonsignificant in both contrasts; (B) events that were significant in the W environment contrast but nonsignificant in the WL environment contrast; (C) conversely, events that were nonsignificant in the W environment contrast but significant in the WL environment contrast; and (D) events that were significant in both contrasts. Scatterplots in panels E-H show the difference in inclusion level for each alternative splicing event between W and WL individuals from a crowded maternal environment (on the x-axis) versus from an uncrowded maternal environment (on the y-axis) for four categories: (E) events that were nonsignificant in both contrasts; (F) events that were significant in the crowded environment morph contrast but nonsignificant in the uncrowded environment morph contrast; (G) conversely, events that were nonsignificant in the crowded environment morph contrast but significant in the uncrowded environment morph contrast; and (H) events that were significant in both contrasts.

In comparison, the two morph-based contrasts (between W and WL individuals for each of the two maternal environments; Figure 2E-H) shared more commonalities. Around 50% (crowded maternal environment: 55.9%; uncrowded: 48.6%) of the events significant in one morph contrast were also significant in the other morph contrast (Figure 2H), while ∼50% were exclusive to the contrast in question (Figure 2F-G). The rate of shared significant events was higher than chance (*X*^2^, p < 2.2×10^-16^). Events that were significant in both morph contrasts almost always (97.8%) had the same direction of effect in both, across a wide range of inclusion level values (Figure 2H, upper right and lower left quadrants; more often than would be expected due to chance, *X*^2^, p < 2.2×10^-16^). When events were nonsignificant in one or both contrasts, we found that they still tended to share the same direction of effect more often than would be expected due to chance across all significance categories (*X*^2^; both contrasts nonsignificant, Figure 2E, p = 1.27×10^-07^; crowded significant and uncrowded nonsignificant, Figure 2F, p = 3.63×10^-12^; crowded nonsignificant and uncrowded significant, Figure 2G, p < 2.2×10^-16^).

### 3.4. Effects of morph and environment may be confounded in some sampling schemes

Many polyphenic systems only produce each morph in their respective environments, meaning that environment is always confounded with morph in these systems. We examined the consequences of confounding morph identity and early life environment by comparing W individuals from the crowded maternal environment to WL individuals from the uncrowded maternal environment, matching each maternal environment with the morph it favors. We found 1,543 significant alternative splicing events in 929 genes (Figure 3A, Table 1).

**Figure 3.**
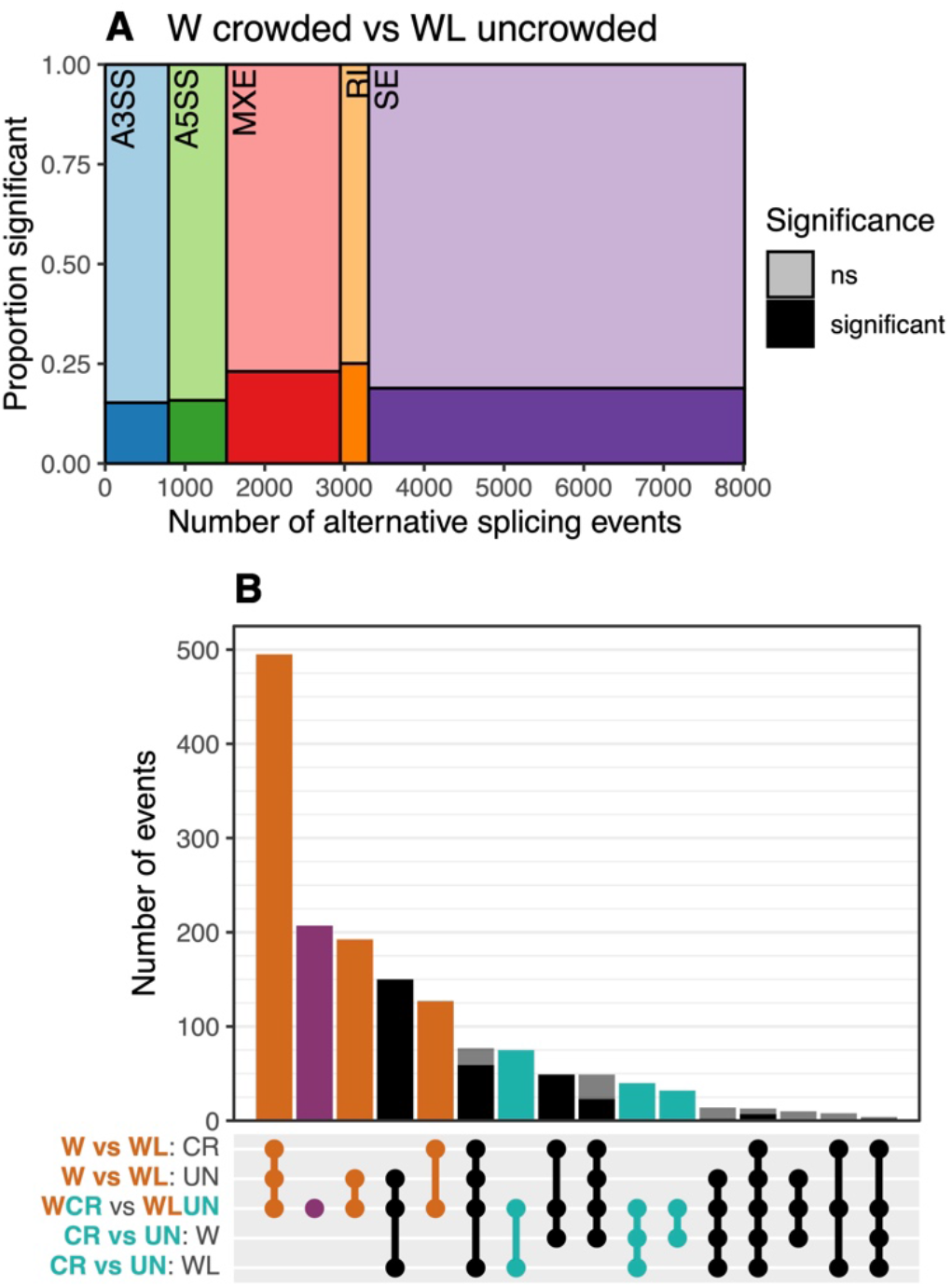
The effect of confounding environment with morph effects. (A) The number of significant alternative splicing events of each splicing type in the W/crowded versus WL/uncrowded paired contrast is shown in a mosaic plot. The x-axis shows the number of total events that were examined for each splicing type (each represented with a different color), while the y-axis shows the number/proportion of those events that were significant (FDR < 0.05) (significant: dark shade; nonsignificant: light shade). (B) The number of significant (FDR < 0.05) events that are shared between the W/crowded versus WL/uncrowded (WCR vs WLUN) contrast and one or more of the other four paired contrasts is shown in an upset plot. The x-axis shows sets of paired contrasts; the contrasts included in a set are represented by dots connected by a line. The y-axis shows the number of events that were found significant in all of the contrasts included in a given set. Dark/intense shades (i.e., black, orange, purple, teal) show events which were significant and had the same direction in all contrasts included in a set, while pale/transparent shades (i.e., gray) show events which did not have the same direction in all contrasts included in a set.

To assess the contributions of environment and morph identity when these factors are confounded, we quantified the proportion of these events that were shared with the other paired contrasts which isolate a single factor (environment or morph; Figure 2). The largest fraction was shared with the other two morph-type contrasts (Figure 3B), and most of the events shared between the two single-environment morph contrasts were also significant in the W/crowded versus WL/uncrowded contrast (634 out of 712 events; Figure S4), suggesting that an experimental design that confounds morph and environment can succeed fairly well at detecting essential morph differences. However, the next-largest category of events (269 events) were those unique to the W/crowded versus WL/uncrowded contrast, and not shared with any other contrast. This, and the presence of some events otherwise only found significant in environment-type contrasts, demonstrate that idiosyncratic, direct effects of the environment are also observed in this confounded experimental design; not all detected differences between morphs are true coordinated morph differences.

### 3.5. Persistent effect of the environment on gene expression

We assessed whether the patterns observed here are specific to alternative splicing by expanding our analysis to a different transcriptomic regulatory level – gene expression. We repeated these analyses and comparisons with gene expression data from the same RNAseq libraries, examining log_2_ fold change in place of inclusion level difference. Differentially expressed genes are largely distinct from differentially alternatively spliced genes (Figure S5), as other studies and systems have also reported (e.g., Grantham & Brisson, 2018; Rogers et al., 2021). However, the patterns of expression differences between groups are broadly similar to the patterns we observe at the level of alternative splicing. Environment-type contrasts produce fewer DEGs than morph-type contrasts (about one-fourth as many; Table S1). When comparing the two environment-based contrasts (Figure S6A-D), we observed little similarity, with only about 5-7% of the DEGs from one contrast also found significant in the other contrast (Figure S6D) and the vast majority exclusive to one contrast or the other (Figure S6B-C). When comparing the two morph-based contrasts (Figure S6E-H), we observed a moderate level of similarity, with 43-66% of the DEGs from one contrast also found significant in the other contrast (Figure S6H) and the other 33-57% exclusive to the contrast in question (Figure S6F-G). We also retested the effect of confounding morph and environment at the level of gene expression; this analysis produced largely similar results (Figure S7).

## 4. Discussion

In this study, we show that the early life environment has long-lasting effects on patterns of alternative splicing in adulthood. This lasting effect goes beyond polyphenic morph identity, with notable differences in alternative splicing between individuals of the same morph whose mothers experienced different environments while their embryos were developing inside them. While splicing differences between W and WL individuals tend to be fairly consistent regardless of maternal environment, the splicing differences that result from the maternal environment are largely dissimilar in W versus WL individuals, indicative of highly distinct morph transcriptomes that are differently perturbed by the effect of the maternal environment. Notably, we were able to detect these effects even though our samples were derived from whole bodies, meaning that differences across tissue types were necessarily summarized and any underlying tissue-specific heterogeneity was likely obscured.

We used expected morph differences in splicing (Grantham & Brisson, 2018) as a scale by which to gauge the magnitude of splicing differences induced by the early life environment. While the morph differences in splicing are certainly the predominant factor in our dataset, the environmental differences are notably large – one-fourth to one-half the size of the morph differences (by simple number of significant differential alternative splicing events, as in Figure 1B-E). This comparison of magnitude is especially valuable since the pea aphid W and WL morphs have dramatically different body plans, easily distinguished with the naked eye, with different tissues and structures including the presence/absence of wings and wing musculature, and sensory and reproductive system differences (Braendle, Davis, Brisson, & Stern, 2006). In contrast, individuals of the same morph from different maternal environments are indistinguishable by eye. In light of the physical differences between morphs versus between individuals of the same morph from different environments, the relative magnitude of alternative splicing differences induced by the environment is particularly striking.

The pea aphid morphs are highly organized and almost certainly have adaptive relevance (e.g., Braendle et al., 2006; Deem, Gregory, Liu, Ziabari, & Brisson, 2024), as is generally assumed to be the case for polyphenic systems (e.g., Simpson et al., 2011). While we observe that the early life environment has a large effect on patterns of alternative splicing, we speculate that this effect may be less organized than that of morph, with uncertain consequences for fitness. We note that the environmental signal in this case has relevance to stress. High conspecific density might be more simply termed ‘crowding stress’, and other factors that may also trigger the wing polyphenism, such as poor host plant quality (e.g., Muller, Williams, & Hardie, 2001; Purandare, Tenhumberg, & Brisson, 2014) likewise are stress-relevant. Changes in alternative splicing in response to environmental stress can be associated with both stress-tolerant and stress-intolerant phenotypes (e.g., Cheng et al., 2025; Tan et al., 2019), with different patterns, and associated consequences range from cognitive dysfunction in rats (Schwarz et al., 2018) to beneficial contributions to pathogen resistance and drought tolerance in plants (Cheng et al., 2025; Mastrangelo, Marone, Laidò, De Leonardis, & De Vita, 2012) and heat tolerance in fish (Tan et al., 2019). Therefore, while the effect of the environment on alternative splicing that we observe here is not as highly organized as the morph determination system, various components of the environmental response could be organized or noisy, and could have beneficial or disadvantageous consequences.

Since pea aphids reproduce clonally, our work provides an especially clear example of the effects of the environment, with no genetic contribution to the differences between individuals. As there are no genetic differences (barring spontaneous mutations), and the response to the environment is long-lasting and not merely immediate, the mechanism must be epigenetic.

Epigenetic markers have long been known to be environmentally responsive, and responsible for persistent environmental effects in many taxa (e.g., Aubin-Horth & Renn, 2009; Champagne, 2010). Mechanistically, splicing is strongly impacted by chromatin structure (Kossowska, 2021) and alternative splicing is closely connected to epigenetic marks (Do, Shanak, Barghash, & Helms, 2023; Hu, Greene, & Heller, 2020), especially in the case of skipped exons for some taxa (Huang et al., 2024). Thus, epigenetic mechanisms almost certainly underlie the differences in alternative splicing that we observe between the genetically identical aphids in our study, and this is a promising avenue for further study.

We note that the striking effects of the early life environment which we demonstrate here may be a complicating factor in many studies of polyphenic systems. This study carefully delineates the distinct signatures of polyphenic morph and of maternal environment in pea aphids. This is made possible by the presence of a small number of individuals of the non-favored morph being produced in each maternal environment. Our results show that in an experimental design that confounds these factors, a variety of effects emerge, including differences that we suggest are central to morph identity, as well as differences stemming directly from the environment with no evident relation to morph. This is an important consideration for studies of morph development and function, as environment and morph effects will generally be confounded in natural populations most or all of the time, but we show that these are distinguishable as two separate effects which are better studied separately when possible.

The early life environment is broadly relevant to health and psychology, where epigenetic mechanisms are increasingly implicated (e.g., Godfrey, Costello, & Lillycrop, 2015; Szyf, 2009). Recently, alternative splicing has also been linked to psychological phenotypes, including addiction (Bhatnagar & Heller, 2025) and the response to social stress (Reshetnikov, Kisaretova, Ershov, Merkulova, & Bondar, 2021; F. R. Wang, Yang, Ren, Chen, & Liu, 2023). Our findings, demonstrating long-lasting splicing effects of early life stress-related cues in a simple insect model, add to this growing body of work.

This study’s outlook begins to bridge the gap between studies of polyphenism and work on health-related effects of the early life environment. Several features of the pea aphid system encourage us to draw parallels. First, pea aphid embryos develop within the mother and experience the environment by way of the mother, a system with similarities to mammalian pregnancy. Second, pea aphids do not undergo pupation or metamorphosis, but gradually mature to adulthood. This simplifies comparisons between aphids and mammalian systems. At the same time, aphids have the advantage that genetically identical individuals may be easily obtained, improving our power to detect and differentiate the striking effect of the early life environment.

## Conclusions

This study reveals striking, long-lasting effects of the early life environment on patterns of alternative splicing in the pea aphid, *Acyrthosiphon pisum*. We carefully distinguish two classes of effects of the early life environment on the adult transcriptome: organized effects of polyphenic morph, and direct effects of the early life environment which are unrelated to morph identity. The direct effects of the early life environment are unexpectedly large in magnitude, nearly rivaling that of morph. This study (1) reveals a need for greater attention to direct (non-morph) effects of the early life environment in studies of polyphenism; and (2) connects to overarching efforts to understand the significance of early life experiences in both biology and medicine.

## Acknowledgements

Lauren Gregory and Julia McDonough generated the RNAseq data employed in this study. We thank Kevin Deem for his helpful input. This work was supported by the National Institute of General Medical Sciences of the National Institutes of Health, USA under award number R35GM144001, and by the University of Rochester.

## Supplemental figures and tables

**Figure S1.**
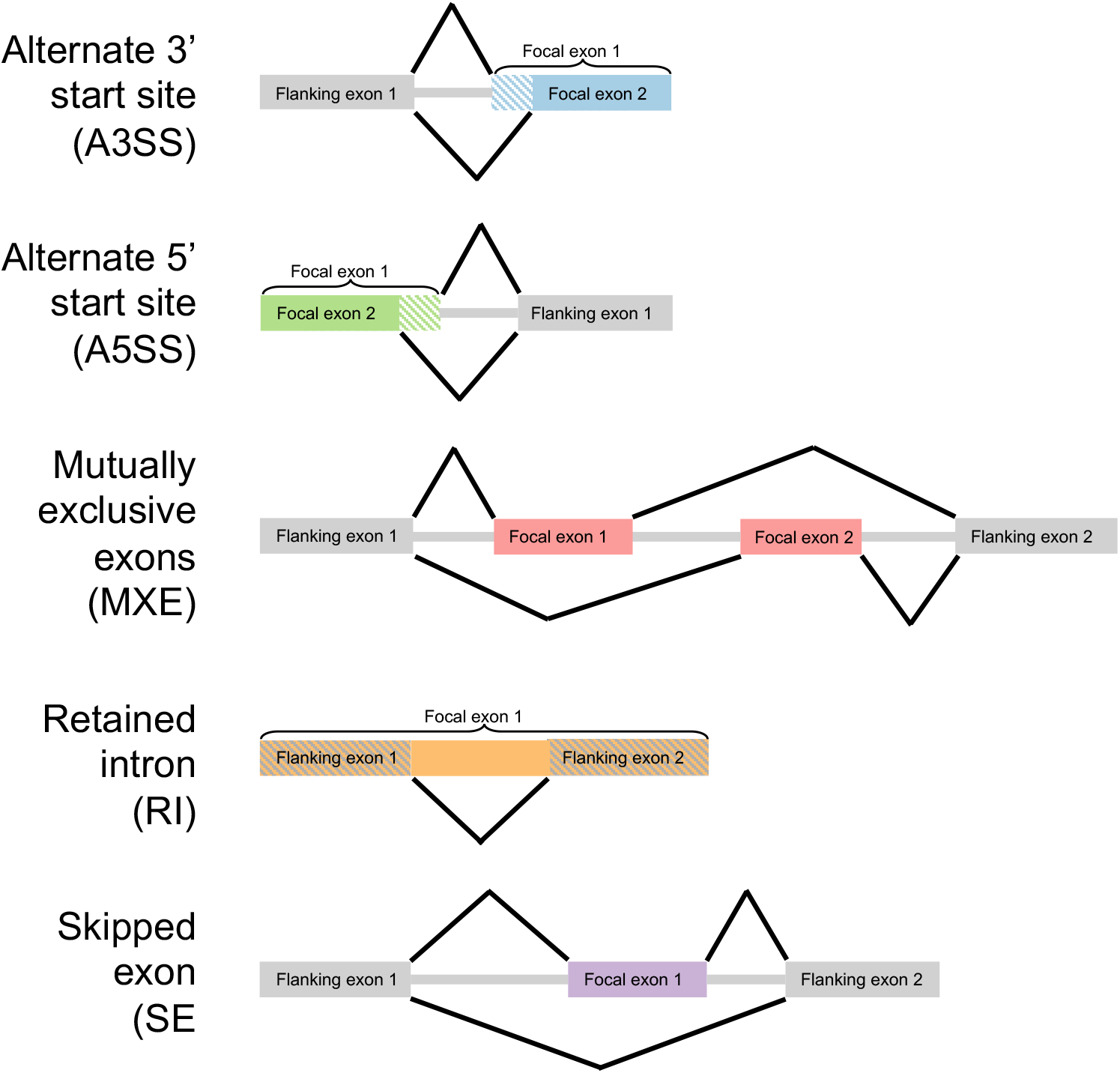
Schematic representations of the five types of alternative splicing events assessed in this work. Boxes (colored for focal exons, gray for flanking exons) represent exons and gray horizontal bars represent introns in each transcript model. White-and-color hatching in A3SS and A5SS shows exon areas that are sometimes included and sometimes excluded, while gray-and-color hatching in RI shows areas that are sometimes assigned to the focal exon and other times assigned to the flanking exons. Black angled lines show exon junctions by connecting exons to be spliced together, bracketing regions that are removed from the transcript. Two splice forms were assessed for each event type: an “inclusion form” indicated by the exon junctions shown above the transcript model and a “skipping form” indicated by the exon junctions shown below the transcript model. In A3SS, A5SS, and SE events, the “inclusion form” is the longer form that includes the focal exon or additional focal exon section. In MXE events, the “inclusion form” is the form that includes the first of the two exons to be encountered when traveling in the direction of transcription, and the “skipping form” is the form that includes the second of the two exons to be encountered, regardless of length. In RI events, the “inclusion form” is again the longer form and includes the focal section without employing any splice junctions, while the “skipping form” splices out a central section between two flanking exons.

**Figure S2.**
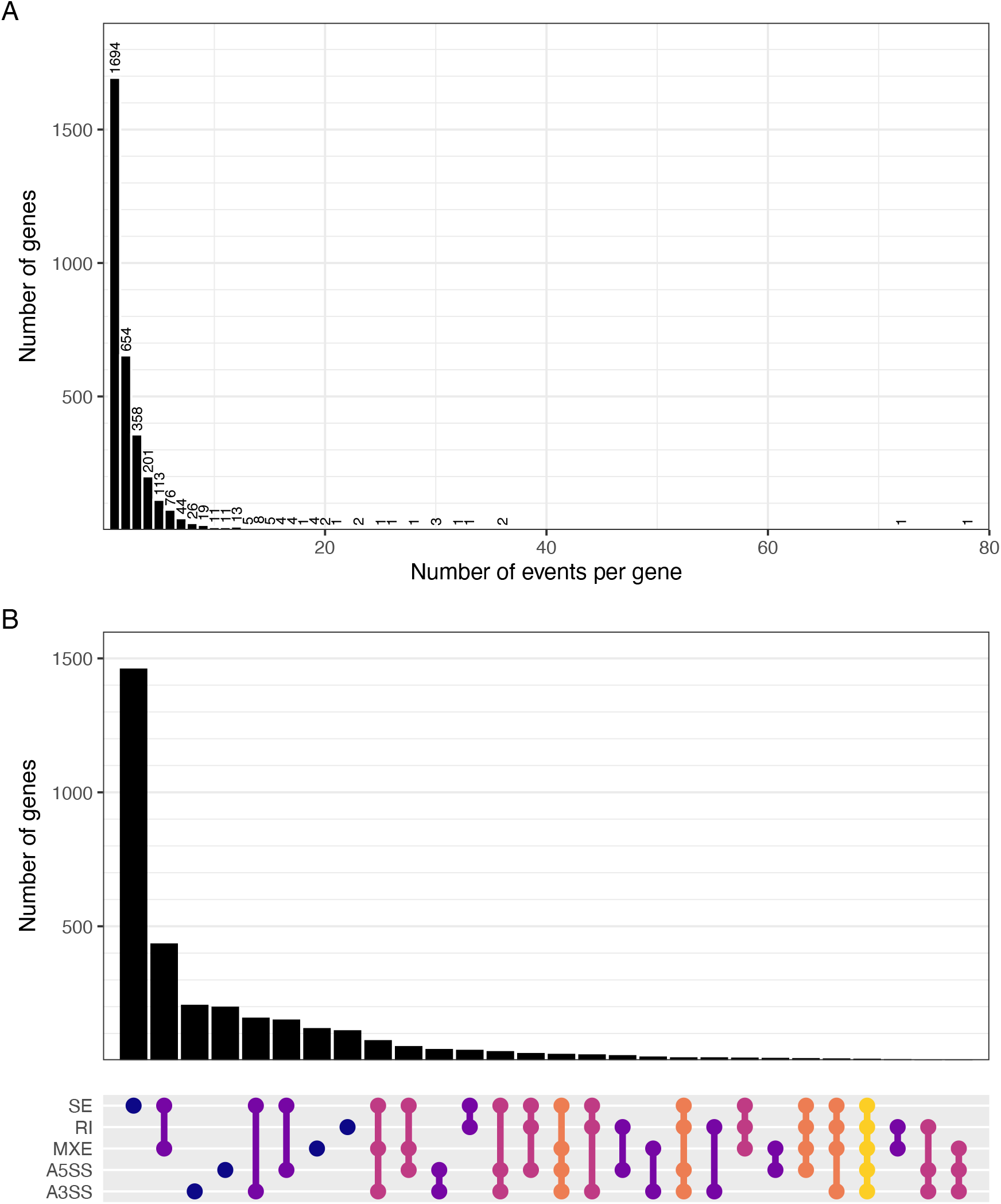
(A) Histogram showing the number of genes containing one or more alternative splicing events included in the analysis post-filtering. The x-axis shows the number of events per gene and the y-axis shows the number of genes with a given number of events. (B) Upset plot showing the number of genes that contain different combinations of alternative splicing event types: skipped exon (SE), retained intron (RI), mutually exclusive exons (MXE), alternative 5’ start site (A5SS), and alternative 3’ start site (A3SS). The x-axis shows sets of alternative splicing event types; the types included in a set are represented by dots connected by a line. The y-axis shows the number of genes that contained the splice types included in a given set.

**Figure S3.**
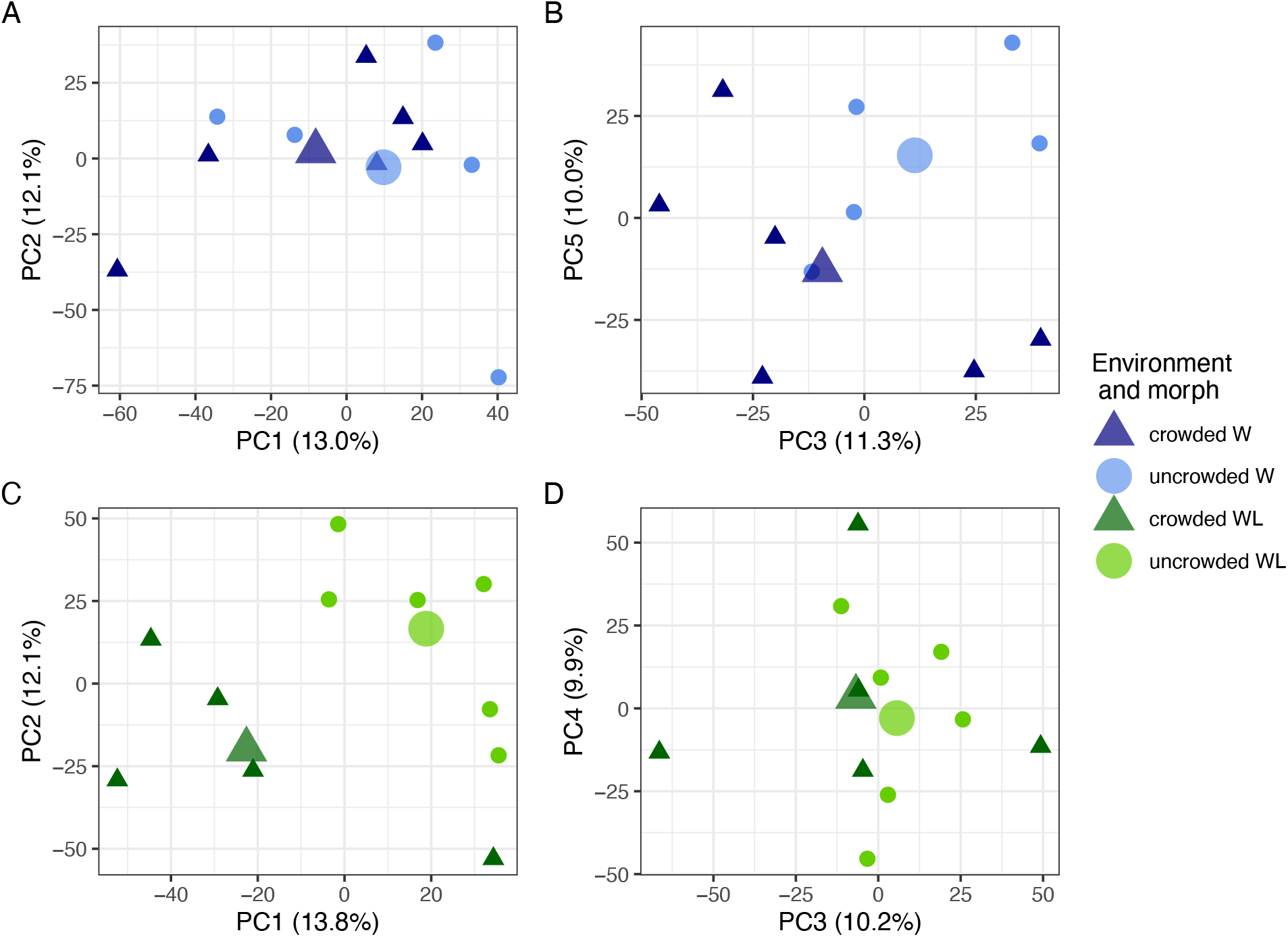
Principal component analysis (PCA) of per-sample inclusion level values for 8016 alternative splicing events conducted separately for winged (W) individuals, in blue (A-B), and wingless (WL) individuals, in green (C-D), reveals the effect of environment on PC3 and PC5 for W individuals (B) and PC1 and PC2 for WL individuals (C). Individual samples from each group are represented by small points (W: blue; WL: green; crowded: triangle, dark shade; uncrowded: circle, light shade), while large points represent the group means.

**Figure S4.**
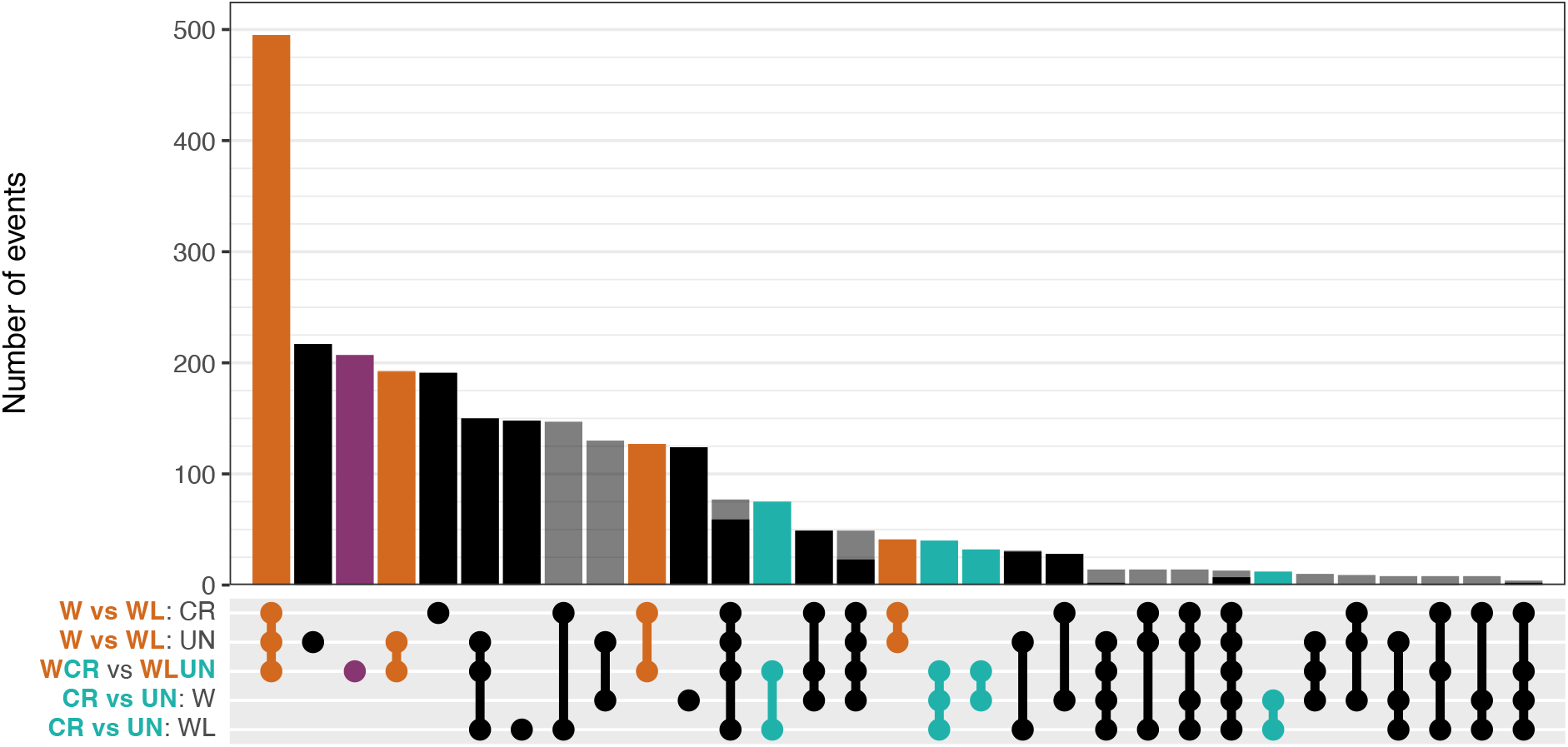
The number of significant (FDR < 0.05) events that are shared between any of the five paired contrasts, or only found in a single paired contrast, is shown in an upset plot. The x-axis shows sets of paired contrasts; the contrasts included in a set are represented by dots connected by a line. The y-axis shows the number of events that were found significant in all of the contrasts included in a given set. Dark/intense shades (i.e., black, orange, purple, teal) show events which were significant and had the same direction in all contrasts included in a set, while pale/transparent shades (i.e., gray) show events which did not have the same direction in all contrasts included in a set.

**Figure S5.**
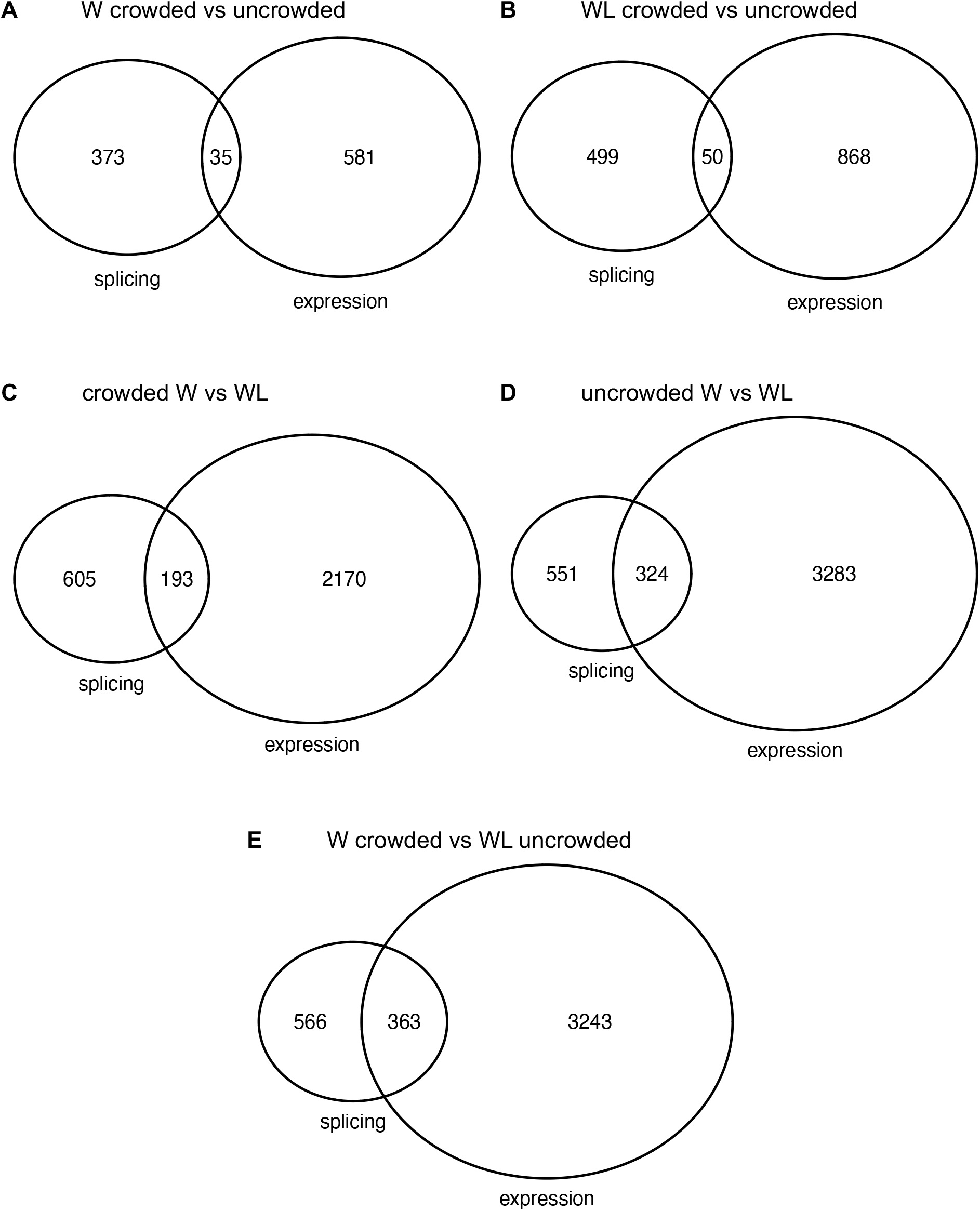
The populations of genes exhibiting significant differential alternative splicing (“splicing”) and genes exhibiting significant differential expression (“expression”) are compared with a Venn diagram for each of the five paired contrasts to show the number of genes exhibiting only one or the other, and the number of genes exhibiting both.

**Figure S6.**
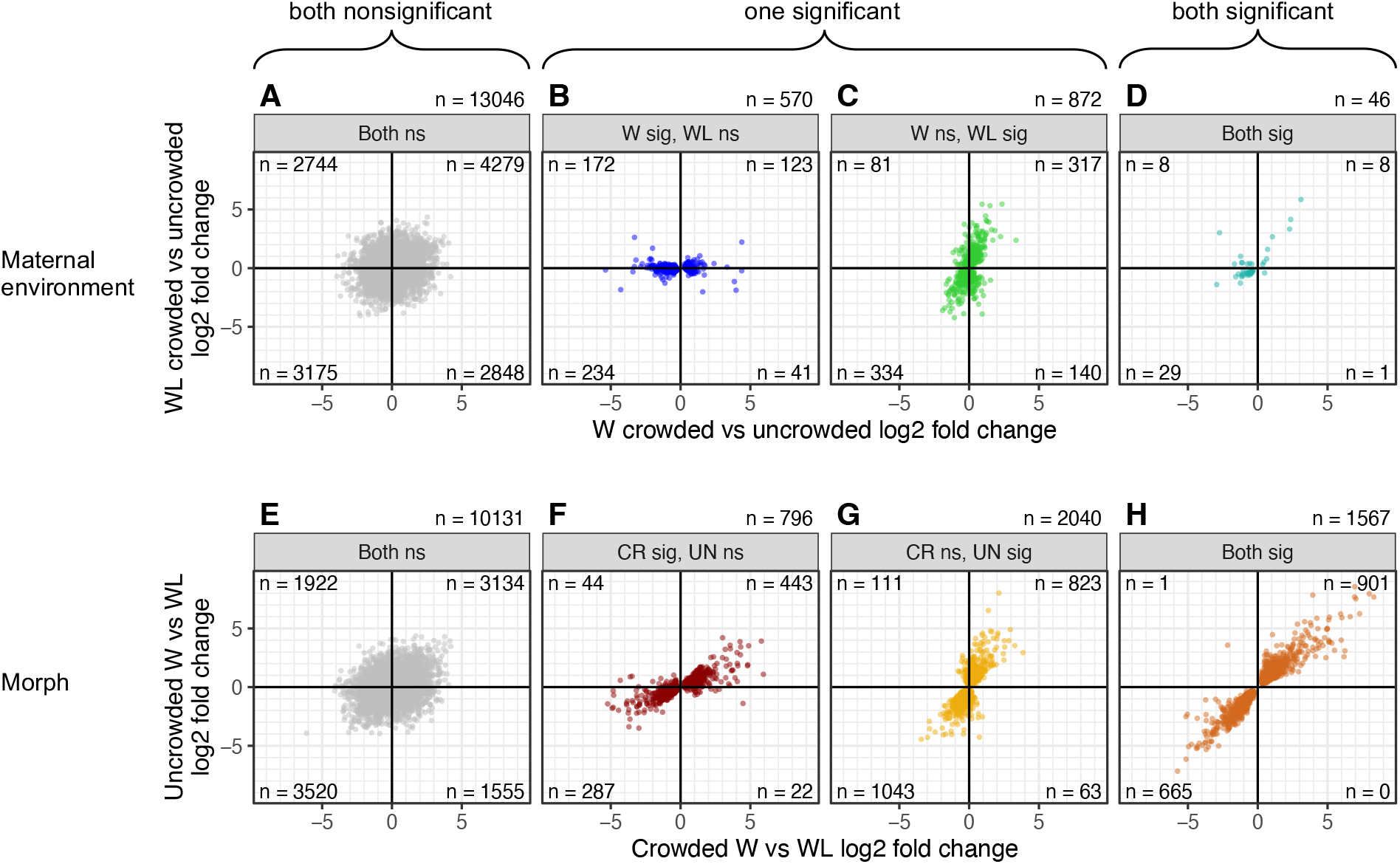
The consistency of effects of morph and environment were assessed at the level of gene expression. (A-D) Scatterplots show the difference in log_2_ fold change for each gene between individuals from a crowded maternal environment versus from an uncrowded maternal environment for W (on the x-axis) versus WL (on the y-axis) for four significance categories; see Figure 2A-D for the alternative splicing version of this analysis. (E-H) Scatterplots show the difference in log_2_ fold change for each gene between W and WL individuals from a crowded maternal environment (on the x-axis) versus from an uncrowded maternal environment (on the y-axis) for four significance categories; see Figure 2E-H for the alternative splicing version of this analysis.

**Figure S7.**
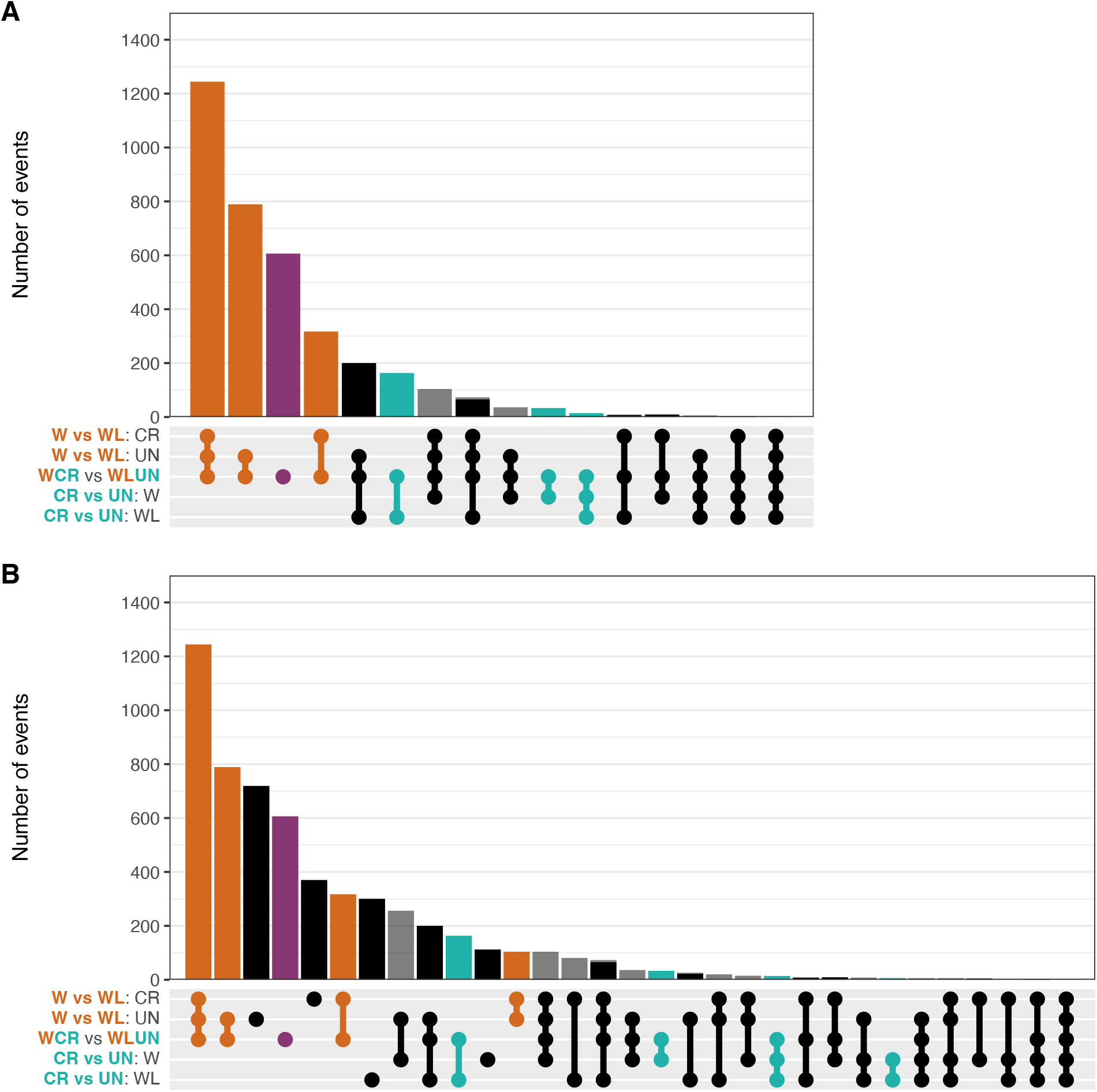
The number of significant DEGs that are shared between any of the five paired contrasts was assessed. (A) The number of significant (FDR < 0.05) DEGs that are shared between the W/crowded versus WL/uncrowded (WCR vs WLUN) contrast and the other four paired contrasts is shown in an upset plot; see Figure 3B for the alternative splicing version of this analysis. The x-axis shows sets of paired contrasts; the contrasts included in a set are represented by dots connected by a line. The y-axis shows the number of events that were found significant in all of the contrasts included in a given set. Dark/intense shades (i.e., black, orange, purple, teal) show events which were significant and had the same direction in all contrasts included in a set, while pale/transparent shades (i.e., gray) show events which did not have the same direction in all contrasts included in a set. (B) The number of significant (FDR < 0.05) DEGs that are shared between any of the five paired contrasts, or only found in a single paired contrast, is shown in an upset plot as in (A); see Figure S5 for the alternative splicing version of this analysis.

---

**Table S1.** The number of genes examined in the gene expression analyses in this study after filtering (see Methods), and the number of significant DEGs for each of five paired contrasts. W CR vs UN, winged crowded vs winged uncrowded; WL CR vs UN, wingless crowded vs wingless uncrowded; CR W vs WL, winged crowded vs wingless crowded; UN W vs WL, winged uncrowded vs wingless uncrowded; W CR vs WL UN, winged crowded vs wingless uncrowded.

|  | # of genes |
| --- | --- |
| All genes being considered | 14534 |
| W CR vs UN significant DEGs | 616 |
| WL CR vs UN significant DEGs | 918 |
| CR W vs WL significant DEGs | 2363 |
| UN W vs WL significant DEGs | 3607 |
| W CR vs WL UN significant DEGs | 3606 |

**Table S2.** Significant GO terms from a gene ontology (GO) analysis conducted on the list of genes containing significant differential alternative splicing events for each of five paired contrasts: WCRUN, winged crowded vs winged uncrowded; WLCRUN, wingless crowded vs wingless uncrowded; CRWWL, winged crowded vs wingless crowded; UNWWL, winged uncrowded vs wingless uncrowded; WCRWLUN, winged crowded vs wingless uncrowded. GO terms are categorized as biological process (BP), cellular component (CC), or molecular function (MF) terms. GO terms that were significant in at least one of the paired contrasts are included in the table. The p-value for each term in each of the five contrasts is reported (NA if the GO analysis for a given contrast did not return a p-value for a given term). Significant (p<0.05) p-values are marked with *.

| Category | Term | WCRUN<br>p-value | WLCRUN<br>p-value | CRWWL<br>p-value | UNWWL<br>p-value | WCRWLUN<br>p-value |
| --- | --- | --- | --- | --- | --- | --- |
| BP | cellular response to amino acid starvation | 0.00581* | 0.749 | 1 | 0.098 | 0.653 |
| BP | anatomical structure morphogenesis | 0.00741* | 0.493 | 0.579 | 0.848 | 0.552 |
| BP | muscle contraction | 0.0278* | 0.694 | 0.462 | 0.214 | 0.562 |
| BP | positive regulation of striated muscle contraction | 0.0416* | 0.447 | 0.154 | 0.188 | 0.206 |
| BP | positive regulation of sarcomere organization | 0.0416* | 0.447 | 0.154 | 0.188 | 0.206 |
| BP | nervous system development | 0.164 | 0.00141* | 0.0515 | 0.0986 | 0.0247* |
| BP | potassium ion transmembrane transport | 0.794 | 0.0111* | 0.154 | 0.98 | 0.906 |
| BP | intracellular signal transduction | 0.603 | 0.0112* | 0.855 | 0.374 | 0.218 |
| BP | regulation of DNA-templated transcription | 0.262 | 0.0179* | 0.331 | 0.0468* | 0.286 |
| BP | mRNA splice site recognition | 1 | 0.0198* | 0.0497* | 0.0673 | 0.0774 |
| BP | regulation of RNA splicing | 0.602 | 0.0225* | 0.0124* | 0.302 | 0.334 |
| BP | signal transduction | 0.482 | 0.0344* | 0.784 | 0.149 | 0.413 |
| BP | regulation of developmental process | 0.651 | 0.0387* | 0.317 | 0.39 | 0.728 |
| BP | animal organ development | 0.952 | 0.0397* | 0.95 | 0.463 | 0.703 |
| BP | regulation of alternative mRNA splicing, via spliceosome | 0.117 | 0.0429* | 0.364 | 0.424 | 0.152 |
| BP | RNA splicing | 0.842 | 0.0883 | 0.0365* | 0.165 | 0.198 |
| BP | axon guidance | 1 | 0.158 | 0.426 | 0.0977 | 0.021* |
| CC | lysosome | 0.0421* | 0.523 | 0.359 | 0.657 | 0.246 |
| CC | adherens junction | 0.0769 | 0.0103* | 0.0285* | 0.128 | 0.00853* |
| CC | membrane | 0.914 | 0.0169* | 0.383 | 0.828 | 0.12 |
| CC | neuron projection | 0.402 | 0.0185* | 0.455 | 0.34 | 0.807 |
| CC | postsynaptic density | 1 | 0.0187* | 0.476 | 0.556 | 0.599 |
| CC | plasma membrane | 0.0774 | 0.0277* | 0.0651 | 0.0547 | 0.00662* |
| CC | dendritic spine | NA | 0.0403* | 0.764 | 0.426 | 0.824 |
| CC | neuronal cell body | 1 | 0.0435* | 0.969 | 0.219 | 0.487 |
| CC | endosome | 0.705 | 0.432 | 0.0285* | 0.0535 | 0.5 |
| CC | cytoplasm | 0.186 | 0.311 | 0.732 | 0.0365* | 0.877 |
| MF | signaling adaptor activity | 0.0445* | 1 | 0.574 | 1 | 0.647 |
| MF | protein binding | 0.0448* | 0.00983* | 0.0229* | 0.208 | 0.108 |
| MF | RNA binding | 0.482 | 0.0288* | 0.0154* | 0.218 | 0.23 |
| MF | tau-protein kinase activity | 1 | 0.0391* | 0.759 | 0.802 | 0.824 |
| MF | structural molecule activity | 0.0705 | 0.147 | 0.00395* | 0.00722* | 0.181 |
| MF | calmodulin binding | 0.16 | 0.582 | 0.0376* | 0.0748 | 0.104 |
| MF | mRNA binding | 0.301 | 0.507 | 0.0489* | 0.195 | 0.388 |
| MF | actin filament binding | 0.114 | 0.0608 | 0.144 | 0.0228* | 0.0183* |
| MF | microtubule binding | 0.968 | 0.834 | 0.142 | 0.124 | 0.0013* |
| MF | cell-cell adhesion mediator activity | NA | 0.0779 | 0.751 | 0.289 | 0.0421* |

